# Layer Va neurons, as major presynaptic partners of corticospinal neurons, play critical roles in skilled movements

**DOI:** 10.1101/2024.06.28.601172

**Authors:** Fumiyasu Imai, Ken Matsuura, Emily Yang, Kelsey Klinefelter, Gabriella Alexandrou, Agustín Letelier, Hirohide Takatani, Fumitaka Osakada, Yutaka Yoshida

**Author notes:** Corresponding author and lead contact Yutaka Yoshida.

## Abstract

Corticospinal neurons (CSNs) are located in the cortex and projecting into the spinal cord. The activation of CSNs, which is associated with skilled motor behaviors, induces the activation of interneurons in the spinal cord. Eventually, motor neuron activation is induced by corticospinal circuits to coordinate muscle activation. Therefore, elucidating how the activation of CSNs in the brain is regulated is necessary for understanding the roles of CSNs in skilled motor behaviors. However, the presynaptic partners of CSNs in the brain remain to be identified. Here, we performed transsynaptic rabies virus-mediated brain-wide mapping to identify presynaptic partners of CSNs (pre-CSNs). We found that pre-CSNs are located in all cortical layers, but major pre-CSNs are located in layer Va. A small population of pre-CSNs are also located outside the cortex, such as in the thalamus. Inactivation of layer Va neurons in *Tlx3-Cre* mice results in deficits in skilled reaching and grasping behaviors, suggesting that, similar to CSNs, layer Va neurons are critical for skilled movements.

Finally, we examined whether the connectivity of CSNs is altered after spinal cord injury (SCI). We found that unlike connections between CNSs and postsynaptic neurons, connections between pre-CSNs and CSNs do not change after SCI.

## Introduction

The brain is highly diverse and is composed of neurons that are classified molecularly, morphologically, physiologically, and functionally^1–6^. In the brain, the cortex has a laminar structure composed of layers I, II/III, IV, Va, Vb, and VI. The cortex also comprises both inhibitory and excitatory neurons, which are subdivided into intratelencephalic tract (IT) and pyramidal tract (PT) neurons. IT neurons project within the telencephalon, while PT neurons project into the spinal cord, as well as the telencephalon, midbrain, and brainstem^10,15^.

Corticospinal (CS) neurons (CSNs), a subset of PT neurons, are located in layer Vb of the sensorimotor cortex and project into the spinal cord. CSNs are involved in voluntary movements, including skilled reaching and grasping behaviors^9,10^. In rodents, CSNs connect to only interneurons in the spinal cord to control skilled movements^11^. Recent studies have identified spinal interneurons that are connected to CSNs ^10,12–14^. For example, CSNs project to Chx10-, vGlut3-, CCK-, Gad2-, and SST-positive interneurons as well as Clarke’s column neurons in the spinal cord^10,12,14,15^. Indeed, ablation of Chx10-, vGlut3-, and Gad2-positive spinal interneurons causes deficits in skilled motor behaviors^10,16^. However, the presynaptic partners of CSNs have not been carefully explored^10,16^.

CS circuits are damaged after spinal cord injury (SCI). Many studies have examined how CS axons are reorganized in the spinal cord after SCI^17^. For instance, loss of PTEN induces axonal regeneration, indicating that mTOR pathways regulate CS regeneration via an intrinsic mechanism^18^. In addition to intrinsic molecules, extrinsic molecules, such as myelin inhibitors, negatively regulate axonal regeneration, and inactivation of these pathways enhances axonal elongation^17^. However, how postsynaptic structures of CSNs are subsequently reorganized and whether connectivity between pre-CSNs and CSNs in the brain is altered remain unclear.

In this study, we performed brain-wide tracing of CSNs in the motor cortex with a rabies virus and found that layer Va neurons constitute the majority of pre-CSNs. Furthermore, inactivating layer Va neurons affects the execution of the pellet-reaching task similarly to inactivating CSNs^10^. Additionally, we performed brain-wide mapping after SCI and found that layer Va neurons constitute the majority of pre-CSNs, suggesting that both layer Va neurons and CSNs are integral to the execution of skilled behavior.

## Results

### Identification of pre-CSNs

To identify presynaptic partners of CSNs, we performed transsynaptic rabies virus tracing. To specifically identify CSNs, a retrograde adeno-associated virus (AAV, AAVrg serotype) was injected into the cervical (C5-6) and lumbar (L4-5) spinal cord (Fig. 1A)^19,20^. CSNs projecting to the cervical spinal cord were located in the rostral forelimb area (RFA) and caudal forelimb area (CFA) of the sensorimotor cortex as well as the secondary sensory area (S2, Fig. 1C). In addition to the cortex, the red nucleus, hypothalamus, pons, and brainstem neurons were also labeled by AAVrg since descending neurons are located in these areas (Fig. S1)^20^. To determine the brain-wide distribution of pre-CSNs, the expression of Cre, TVA (virus envelope protein EvnA receptor), GFP and optimized rabies glycoprotein (oG) was induced by AAVrg-Cre and AAV8-Flex-TVA-P2A-GFP-2A-oG^21,22^. Then, EnvA-coated G-deleted SAD 19 rabies-tdTomato was injected into the RFA, CFA or S2 (Fig. 1D-O)^23^. We first determined the location of the starter neurons expressing both GFP and tdTomato, and we found that they were strictly located at their target sites (Fig. 1G, J, M). Then, neurons expressing only tdTomato were identified as pre-CSNs. The majority of pre-CSNs were observed within the cortex, and small populations were located in the thalamus (Figs. 1E, F). In the cortex, layer V neurons were the main pre-CSNs (40-70%, Fig. 1F). CSNs received inputs mainly from local areas (Figs. 1H, I, K, L, N, O), and more than 60% of the pre-CSNs were located within 1 mm of the starter neurons (76.4 ± 8.2% in the RFA, 66.2 ± 14.7% in the CFA, and 69.7 ± 17.0% in S2). We also found that pre-CSNs were mainly located in contralateral sites, although fewer than 10% (3.4 ± 1.3% in the RFA, 9.4% ± 9.7% in the CFA, and 0.07 ± 0.06% in S2) of the total pre-CSNs were in the cortex.

**Figure 1.**
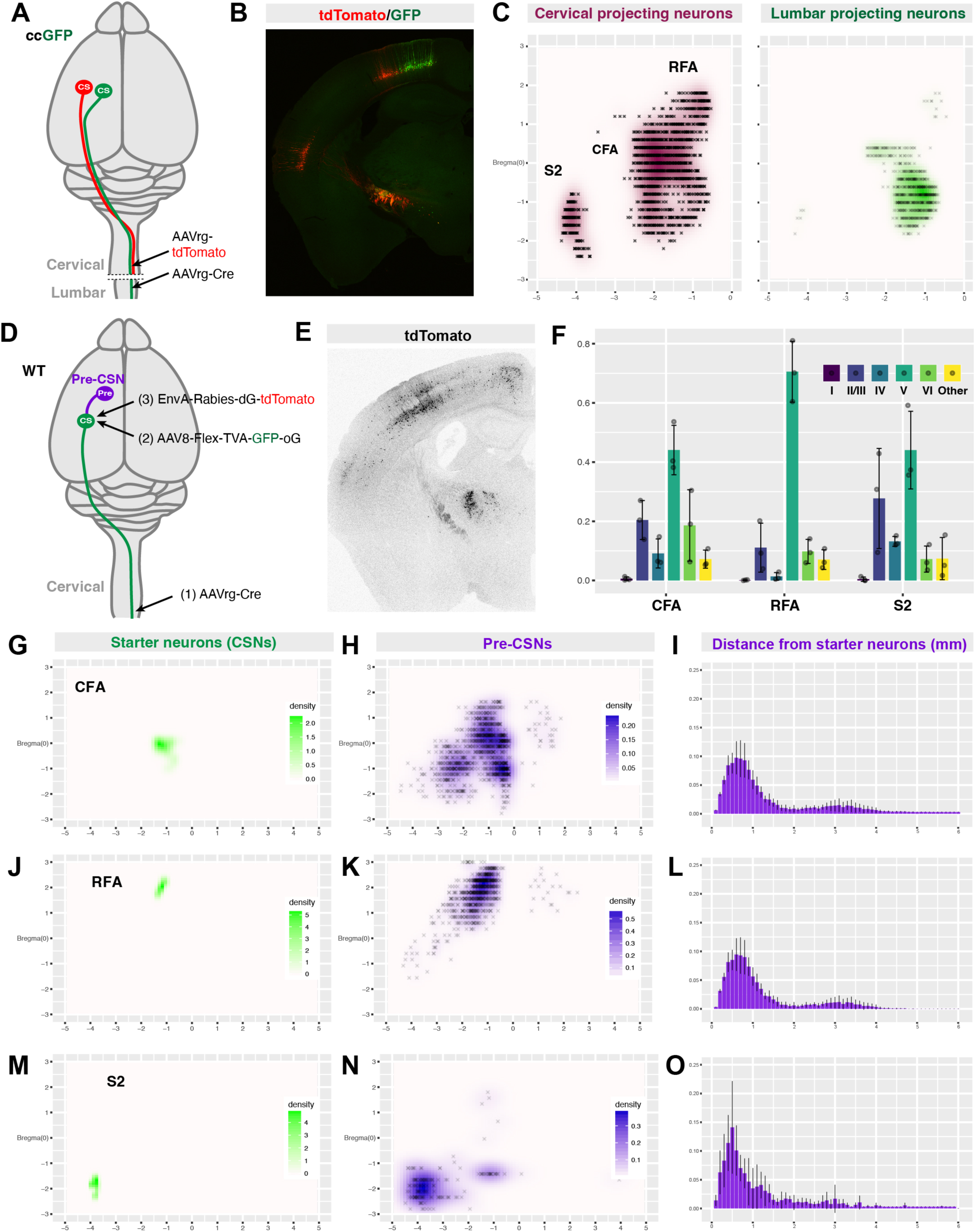
Mapping of pre-CSNs. (A) Schematic drawing of the labeling of CSNs by AAVrg. AAVrg-tdTomato and AAVrg-Cre were injected into the cervical (C6) and lumbar (L5) spinal cords of Cre-ccGFP mice, respectively. (B) Coronal section of the cortex showing tdTomato (cervical)- and GFP (lumbar)-expressing CSNs and hypothalamic neurons. (C) Mapping of cervical CSNs. (D) Schematic drawing of the pre-CSN labeling strategy. TVA and oG expression in CSNs was induced using AAVrg-Cre and AAV8-Flex-TVA-GFP-oG. Then EnvA-rabies-dG-mCherry-infected TVA expression was induced in CSNs. (E) Coronal section of a rabies-dG-mCherry-infected brain. (F) Localization of pre-CSNs. (G-O) Mapping of starter cells (G-I) and pre-CSNs (J-L) and distances between starter cells and pre-CSNs (M-O). AAV8-Flex-TVA-GFP-oG and EnvA-rabies-dG-mCherry were injected into the CFA (G, J, M), RFA (H, K, N) and S2 (I, L, O). RFA: rostral forelimb area, CFA: caudal forelimb area, S2: secondary sensory cortex.

To determine whether the pre-CSNs in layer V were CSNs, we labeled CSNs by injecting CTB into the pons and found that most pre-CSNs in cortical regions were CTB negative (95.4%, Figs. 2A, B), suggesting that these pre-CSNs were not CSNs^24^. Next, to examine whether the pre-CSNs were layer Va neurons, we used Tlx3-Cre mice to visualize layer Va neurons and performed rabies virus tracing using EnvA-coated G-deleted rabies-Flpo and intersectional Flpo and Cre double-dependent AAV-EYFP (Figs. 2C, D)^25,26^. We found that Cre and Flpo double-positive neurons were labeled by the intersectional EFYP viruses, and these neurons projected to the contralateral cortex and striatum, indicating that the pre-CSNs were IT neurons in layer Va (Fig. 2D)^27^. To confirm the projection patterns of pre-CSNs in layer Va, Cre-dependent AAVrg-DIO-Flpo was injected into the contralateral side, and Flpo-dependent AAV8-fDIO-Synaptophysin-mRuby was injected into the ipsilateral cortex of Tlx3-Cre mice (Figs. 2E, F). Although the Synaptophysin-positive neurons were contralaterally projecting layer Va neurons, the majority of presynaptic puncta were located in layer V of the ipsilateral cortex (Fig. 2G). Presynaptic puncta were also observed in the contralateral cortex and the ipsilateral and contralateral striatum (Figs. 2H-K). These data indicate that layer Va neurons constitute the majority of pre-CSNs and that these neurons project to CSNs as well as to the contralateral cortex and striatum.

**Figure 2.**
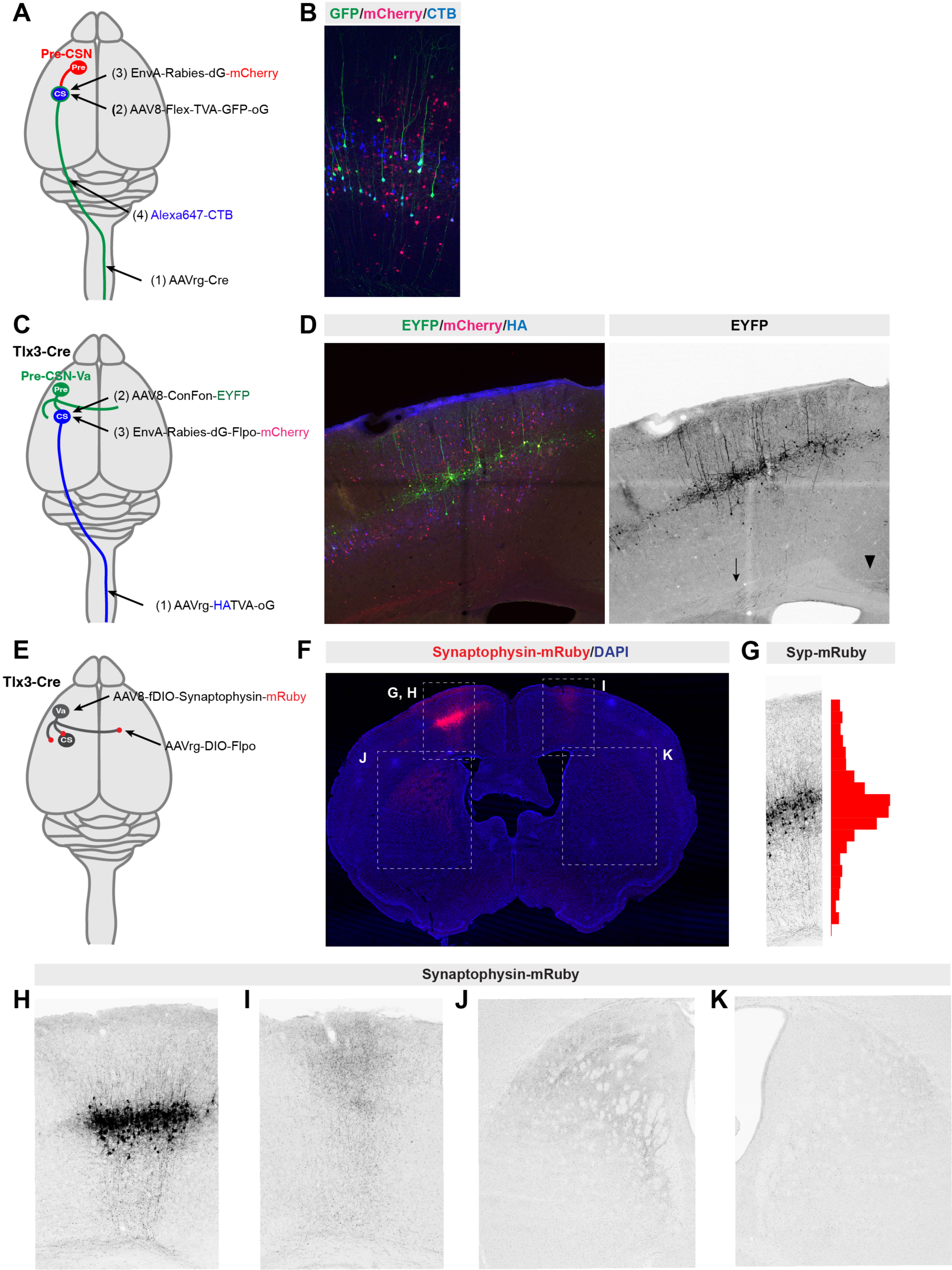
CSNs receive inputs from layer Va neurons. (A, B) Labeling of pre-CSNs and CSNs by rabies viruses (red) and CTB (blue), respectively. Starter cells were marked by GFP (green) and expressed TVA, oG and GFP. (C, D) Pre-CSN-Va neurons were labeled with AAVrg-HATVA-oG, AAV8-Con/Fon-EYFP and EnvA-Rabies-dG-Flpo-mCherry (C). Cortical images of EYFP (green), mCherry (red and black) and HA (blue) (D). (E-K) Labeling of contralaterally projecting layer Va neurons in the contralateral and ipsilateral cortices of Tlx3-Cre mice by AAVrg-DIO-Flpo and AAV8-fDIO-Synaptophysin-mRuby (E). Synaptophysin-mRuby was detected in the ipsilateral cortex (G, H), contralateral cortex (I), ipsilateral cortex (J) and contralateral striatum (K). The density of presynaptic structures was calculated (G, right).

To determine the anatomical connections between layer Va neurons and CSNs, excitatory and inhibitory synapses on the dendrites of CSNs were visualized by injecting AAVrg-PSD95-mCherry and AAVrg-EGFP-Gephyrin, respectively, into the spinal cord (Fig. 3A)^28^. Although small numbers of PSD95-mCherry and EGFP-Gephyrin double-positive structures were observed within cell bodies (arrow in Fig. 3B), the postsynaptic structures of CSNs were found specifically on dendrites (Fig. 3B, C). Both excitatory and inhibitory postsynaptic structures were distributed throughout the cortical layers, but they were mainly located in layer V (EGFP-Gephyrin: 46.9±11.3%, PSD95-mCherry: 61.8±11.7%). In addition to excitatory postsynaptic structures of CSNs, presynaptic terminals of layer Va and layer II/III neurons were labeled with AAV8-Flex-Synaptophysin-EGFP in Tlx3-Cre and SepW1-Cre mice (Fig. 3D-F)^29,30^. The synaptic terminals of layer Va were located mainly in layer V, where the postsynaptic densities of CSNs were located (Fig. 3E). In contrast, SepW1-positive layer II/III terminals were located mainly in layers II/III and V, especially in the upper part of layer V (Fig. 3F). These data indicate that both layer II/III and Va neurons project to CSNs but are spatially segregated.

**Figure 3.**
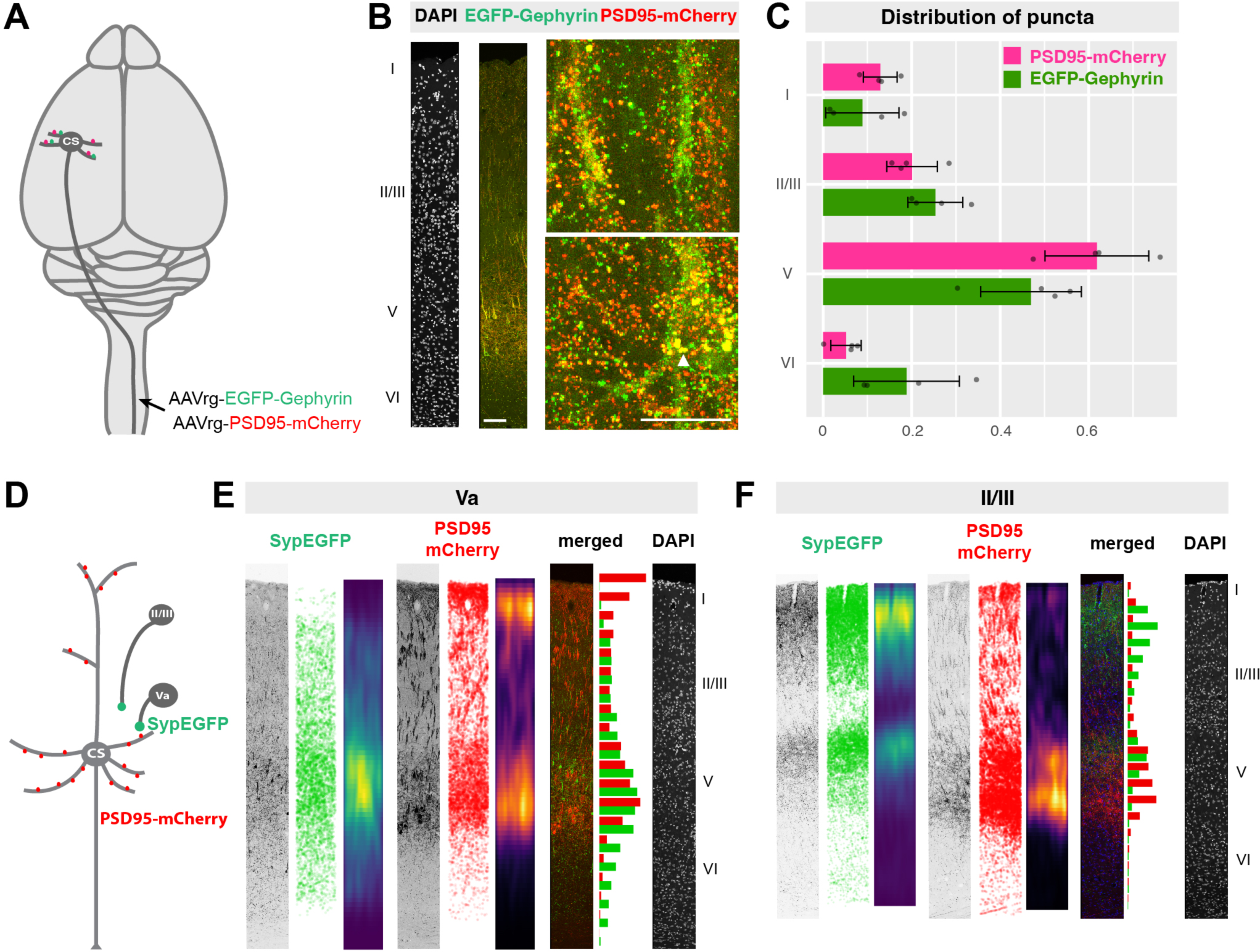
(A-C) Visualization of postsynaptic structures of CSNs by AAVrg-PSD95-mCherry (red) and AAVrg-EGFP-Gephyrin (green) (B, C). AAVrg-PSD95-mCherry (red) and AAVrg-EGFP-Gephyrin (green) were located mainly on layer V (C). (D) Postsynaptic structures of CSNs and presynaptic structures of layer Va neurons were labeled with AAVrg-PSD95-mCherry and AAV8-Flex-SypEGFP, respectively, in Tlx3-Cre mice. (E, F) (left-right) Image of SypEGFP signals, plot of the number of SypEGFP-positive presynaptic structures, plot of the density of presynaptic structures, image of PSD95-mCherry signals, plot of the number of PSD95-mCherry-positive postsynaptic structures, plot of the density of postsynaptic structures, merged image, histogram of the numbers of presynaptic (green) and postsynaptic (red) structures, and image of DAPI staining in Tlx3-Cre (E, Va) and SepW1-Cre (F, II/III) mice.

### Layer Va neurons are critical for skilled behaviors

CSNs are critical for the execution of skilled movements such as reaching and grasping behaviors, and either inactivation or ablation of CNSs causes deficits in pellet-reaching tasks^9,10^. To determine the roles of layer Va neurons in skilled behaviors, we used designer receptors that are exclusively activated by designer drugs (DREADDs)^31^. Inhibitory DREADD viruses (AAV9-Flex-hm4di-mCherry) were injected bilaterally into the sensorimotor cortex of *Tlx3-Cre* mice (Figs. 4A-C)^32^. After training, the success rate increased to 25%. Then, layer Va neurons were inactivated by the administration of deschloroclozaine (DCZ), which is a ligand for the DREADD, to the mice. The success rate decreased, indicating that layer Va neurons play critical roles in reaching behaviors (Fig. 4D, E). To clarify which components were affected during pellet reaching, the failure ratios of reaching, grasping and retrieving were analyzed. The failure ratio did not change, indicating that all the components were affected by inactivation of layer Va neurons in the mice.

**Figure 4.**
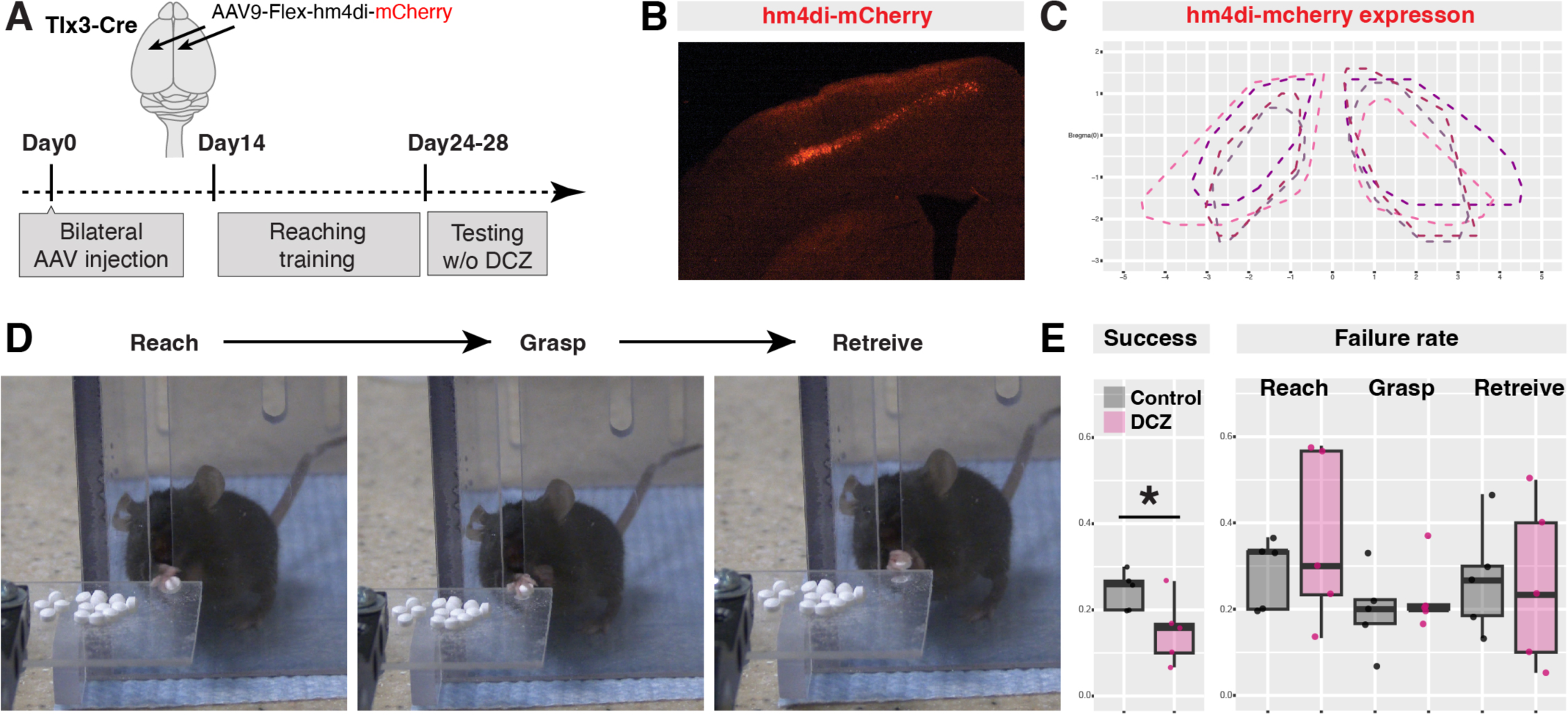
Inactivation of layer Va neurons affects pellet reaching ability. (A) Experimental details of the pellet reaching tests. AAV9-Flex-hm4di-mCherry was injected into the cortex of Tlx3-Cre mice bilaterally, after which the mice were trained on the pellet reaching task. After training, DCZ was administered intraperitoneally. (B) Coronal section of the cortex of an AAV9-Flex-hm4di-mCherry-injected mouse. (C) Mapping of hm4di-mCherry-expressing layer Va neurons. After the reaching test, the brains were sectioned and analyzed. (D) Typical pellet reaching behavior. Mice aimed, started reaching (left, Reach), grasped a pellet (center, Grasp), and then retrieved a pellet before eating it (right, Retrieve). See also the Materials and methods section. (E) Success rate in the presence and absence of DCZ (left) and reaching component analysis (right). The graph shows the failure rate of each component. * p<0.05 (Student’s t test).

### Pre-CSN circuity after SCI

Next, we examined whether connections between layer Va neurons and CSNs are altered after SCI. Although many studies have examined how the axons of CSNs are affected after SCI^17^, a few studies have determined how the dendrites of CSNs are affected after SCI. For example, it has been reported that the spine density of CSNs is reduced in layers II/III and Va after SCI, indicating that the connectivity between pre-CSNs and CSNs changes after SCI^33^. However, this has not yet been tested.

To determine whether the connectivity of CSNs in the brain is altered after SCI, we visualized pre-CSNs and compared them before and after dorsal column lesioning. For this experiment, we used the low-toxicity rabies virus strain CVS-N2c instead of SAD19^34^. We compared the control and SCI groups (Fig. 5A). Unlike in the mice injected with SAD19, many neurons were identified as pre-CSNs in the mice injected with CVS-N2c (3774 ± 689 cells in the CFA after SAD19 injection and 7510 ± 2114 cells in the CFA after CVS-N2c injection). In addition, the distance from the starter neurons was greater after CVS-N2c injection than after SAD19 injection (CVS-N2c: 1.82 ± 0.07 mm, SAD19: 1.01 ± 0.26 mm, Fig. 5B). Furthermore, after SAD19 injection, thalamic neurons were labeled but represented less than 10% of the total pre-CSNs, whereas 20% of the total pre-CSNs were in the thalamus after CVS-N2c injection (Fig. 5C). The majority of pre-CSNs were identified in layer V using CVS-N2c (Fig. 5C, D), similar to what was observed after SAD19 injection. Then, we compared the location of pre-CSNs, including layer Va neurons, before and after SCI, but there were no differences (Fig. 5E). These data suggest that connectivity between pre-CSNs and CSNs is not altered after SCI.

**Figure 5.**
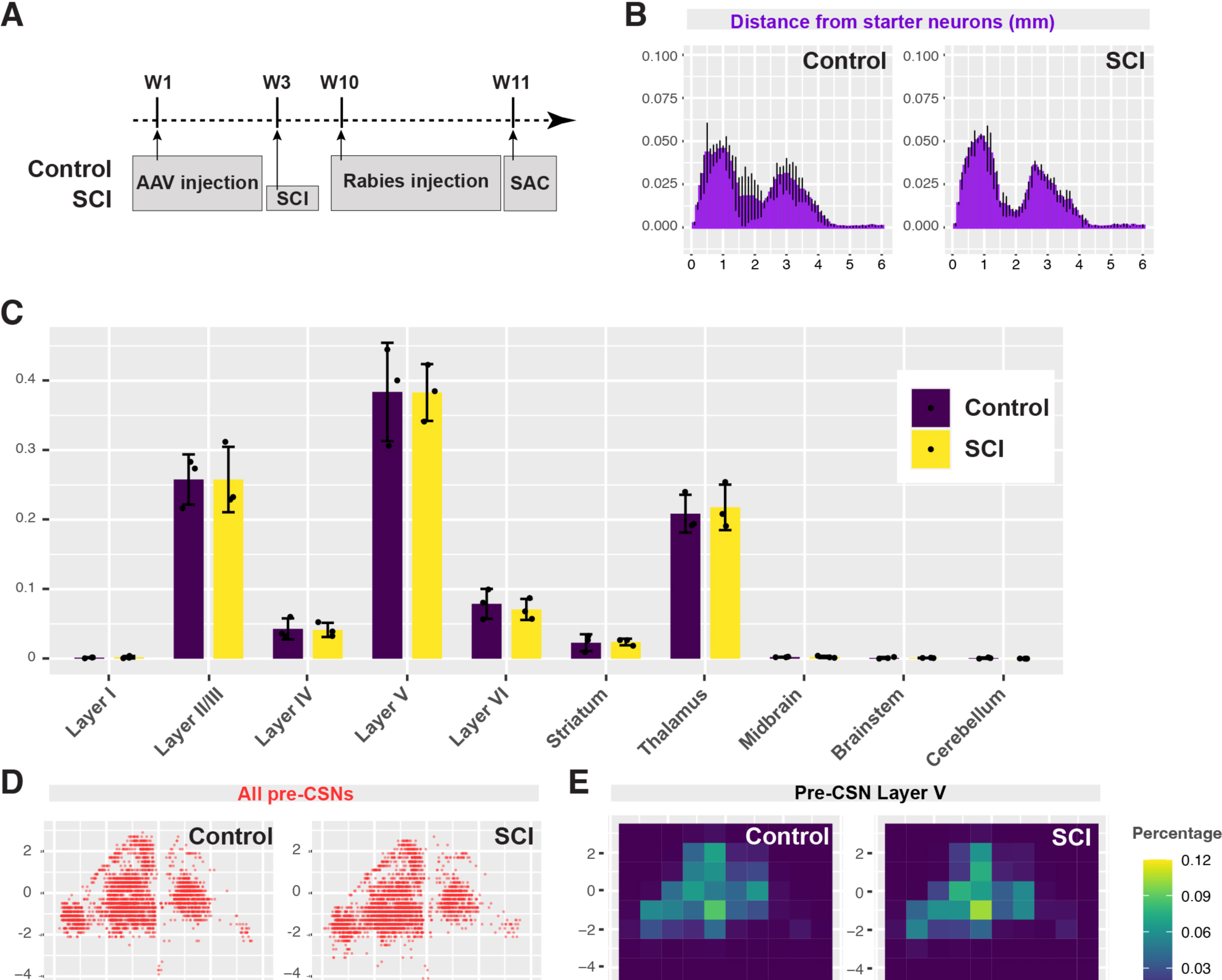
Details of pre-CSN analysis after SCI. (A) TVA and oG expression was induced in CSNs using AAVrg-Cre and AAV8-Flex-TVA-GFP-oG at week 1 (W1). The SCI group received dorsal column lesions at W3. EnvA-rabies-dG-tdTomato was injected at W10, and the mice were sacrificed at W11. (B) Distances between the starter cells and pre-CSNs in the control (left) and SCI (right) groups. (C) Localization of pre-CSNs. (D, E) Mapping of all pre-CSNs (D) and percentage density map of layer V pre-CSN in the control (left) and SCI (right) groups.

## Discussion

Many previous studies have focused on connectivity between CSNs and post-CSNs in the spinal cord^10,12–15^. For example, Chx10-, vGlut3-, and Gad2-positive spinal neurons were identified as post-CSNs, and these interneurons are critical for skilled movements^10,16^. In contrast, little is known about pre-CSNs in the sensorimotor cortex.

In this study, we identified Va neurons as major pre-CSNs in the sensorimotor cortex using trans-synaptic rabies virus tracing with both the SAD19 and CVS-N2c strains^35^. We found that in addition to layer V neurons, layer II/III neurons and thalamic neurons are also pre-CSNs. Similar results were reported for the visual cortex, in which layer V neurons constitute the majority of pre-PT neurons^36,37^. We also found that the inhibition of layer Va neurons affects skilled behavior.

To visualize postsynaptic sites of CSNs, we also labeled excitatory and inhibitory postsynaptic structures with PSD95-mCherry and GFP-gephyrin. The dendrites of CSNs are subdivided into two parts apicobasally, apical dendrites are located mainly in layers I-III and basal dendrites are well branched throughout layers V-VI in the sensorimotor cortex^38^. We found that both PSD95-mCherry and GFP-gephyrin are located mainly in layer V, where the axons of layer Va neurons are located. Indeed, it has been reported that striatum-projecting layer Va neurons also send inputs to CSNs^39^. The number of spines contacting both the apical and basal dendrites of CSNs is increased in after motor training^40^, suggesting that both types of inputs (those projecting to apical dendrites and those projecting to basal dendrites) are changed during training for skilled behaviors^41–44^.

Previous studies have examined how cortical neurons modulate CSN activity^8^. For example, inputs from layers II/III are major inputs to CSNs, and the activity of layer II/III neurons shows a clear correlation with movement^41,45,46^. In addition, layer Va neurons are IT neurons projecting to the contralateral cortex as well as the striatum^4,27,30,47,48^. These neurons also project to CSNs^39^. Contralateral cortex- and striatum-projecting neurons are associated with motor command-like activities, and inactivation of layer Va affects lever task movements^49–51^. We found that inactivation of Va neurons affects performance in the pellet reaching task, indicating that layer Va neurons contribute to skilled motor behaviors. Recent analyses of brain-wide neural activity in individual cortical layers revealed that activation patterns of PlexinD1^On^ IT neurons in layers II/III and Va are different from those of Fezf2^On^ PT neurons in layers V and VI, indicating that brain-wide networks in different cortical layers are critical for skilled behaviors^6^.

To determine whether and how CSN connectivity changes after SCI, we performed brain-wide rabies tracing (Fig. 5). We found that the distribution patterns of pre-CSNs in the cortex and other areas were not altered, and these data indicate that layer Va neurons are major pre-CSNs before and after injury and that the distribution patterns of pre-CSNs are maintained after SCI. However, we cannot exclude the possibility that the single-neuron network changes. To examine pre-CSNs at single-cell resolution, TVA66T, which has low affinity for EnvA and is suitable for local labeling, may be helpful^52^. Recent studies have shown that stimulation of motor neurons, spinal interneurons or CSNs induces motor function recovery after SCI^53–55^. Since layer Va neurons connect with not only CSNs but also other neurons, the combined activation of different neuronal types, such as pre-CSNs (layer Va neurons) and CSNs, may be more efficient than activation of a single neuronal type.

## Materials and methods

### Mice

The following mouse lines were used in this study: C57BL/6J (JAX, #000664), Tlx3-Cre and SepW1-Cre (PL56Gsat, NP39Gsat, MMRC)^25^. Mouse handling protocols and other procedures were approved by the Institutional Animal Care and Use Committee at the Cincinnati Children’s Hospital Research Foundation and Burke Neurological Institute. Both sexes were used for this study.

### Viruses

The pAAV vectors pAAV-hSyn-PSD95-mCherry, pAAV-hSyn-EGFP-Gephyrin, pAAV-hSyn-Flex-SypEGFP and pAAV-fDIO-DTR-mCherry were generated from the original vectors (Addgene #55650, #51509, #73918, #73919, #130909, #124364, and #73919) ^9,26,28,56,57^. The AAV11 capsid gene^58^ was synthesized and inserted into the pAAV2/1 helper vector (Addgene #112862). The following viruses were used in this study: AAVrg-CAG-Cre (UNC viral core), AAV8-CAG-Flex-TVA-P2A-eGFP-2A-oG, EnvA-dG-Rabies-tdTomato (Salk Institute viral core), EnvA-N2C-dG-tdTomato (2×10^9^, Center for Neuroanatomy with Neurotropic Viruses, CNNV)m AAV9-hSyn-DIO-hM4d(Gi)-mCherry (Addgene #44362)^32^, AAVrg-PSD95-mCherry, AAV8-hSyn-Flex-SypEGFP, and AAV8-CAG-DIO-ChRger2-TS-YFP (Addgene #127239) (packaged by Vector Biolabs and Vigene).

### Surgery

We performed viral injection using a stereotaxic frame (Stoelting) and a Nanoject III injector (Drummond). The mice were anesthetized using SomnoSuite (Kent Scientific) with isoflurane (1.5-2%). The following coordinates were used for injections: spinal cord (C5-6 or L5-6): 0.5 mm ML from the midline, 0.5 and 1.0 mm deep from the dorsal surface; RFA: 1.8 mm AP from bregma, 1.5 mm ML from the midline, 0.5 mm deep from the cortical surface; CFA: -0.5 mm AP from bregma, 1.8 mm ML from the midline, 0.5 mm deep from the cortical surface; S2: (-1.8 mm AP from bregma, 3.5 mm ML from the midline, 1.0 mm deep from the cortical surface. For dorsal column lesioning, the cervical spinal cord was exposed, and the C5 dorsal column was lesioned at a depth of 1 mm with Vannas sprig scissors (2.5 mm cutting edge, 150000-08, Fine Science Tools)^59^.

### Behavior

The single-pellet reaching task was carried out as previously reported with some modifications^10,60^. Eight-week-old mice were food restricted so that they maintained 80-90% of their original weight. The reaching chamber was composed of clear plexiglass with 0.5 cm wide slits on the left side, in the center, and on the right side of the front wall. Sucrose pellets (Bio-Serv, #F07595, 20 mg) were placed in front of the slit for the mice to reach, grasp, and retrieve through the slit. During the deprivation/habituation periods (2 days), the mice were placed in the reaching chamber with sucrose pellets for 20 min/day. Subsequently, the mice were trained for shaping (2-7 days). During this period, sucrose pellets were placed around the slits. After shaping training was completed (more than 20 trials per day, greater than 70% of reaches were performed with the dominant hand), the training periods were started on day 7. On day 8, deschloroclozaine (DCZ, MedChemExpress, HY42110, 3 µg/kg) was injected intraperitoneally. All trials on days 7 and 8 were recorded by a video camera (Sony 4K video camera, AX53, 60 fps), and behavior was analyzed manually. The following criteria were used to assess behavior in this study: <u>success:</u> the pellet was grasped, lifted, and eaten; <u>reaching failure</u>: the arm or paw made contact with the pellet but did not grasp it, the underside of the paw/fingers failed to make contact with the pellet, or the paw could not be positioned above the pellet; <u>grasping failure:</u> the paw was positioned above the pellet or the underside of the paw contacted the pellet, but the pellet could not be lifted; <u>retrieval failure:</u> the pellet was grasped and lifted but fell from the paw before being eaten.

### Tissue preparation

The mice were deeply anesthetized with ketamine/xylazine and then perfused with PBS and 4% PFA/PB. After removing the skull and dorsal part of the vertebrae, the tissues were fixed with 4% PFA/PB overnight, and then the brains and spinal cords were separated from the other tissues. The brains and spinal cords were cryoprotected with 30% sucrose/PB two overnights and then embedded in Tissue-Tek OCT compound (Sakura) and cryosectioned (50 µm).

### Immunostaining

The following primary antibodies and reagents were used in this study: rat anti-HA (Roche, 3F10, 1/1000), rabbit anti-GFP (Invitrogen, A-11122, 1/1000), Alexa 647-conjugated donkey anti-rat IgG, Alexa 488-conjugated rabbit (Invitrogen #A48272, #A-21206, 1/1000), and DAPI (Invitrogen, D1306, 5 µg/ml). The sections were mounted with Vectashield (Vector Laboratories).

### Scanning and data analysis

The sections were scanned under SP8 (Leica) and A1RHD25 (Nikon) confocal microscopes. The images were analyzed with IMARIS (Oxford Instruments) and Fiji/ImageJ followed by R and reconstructed using Photoshop and Illustrator (Adobe). For rabies virus mapping, the brains were sectioned at a thickness of 50 µm, and then every 4 sections were scanned. Neurons were counted and reconstructed using R. For synapse counting, individual spots were detected by the IMARIS Spot tool and then reconstructed using R.

## Acknowledgment

The authors would like to acknowledge the Structural and Functional Imaging Core at Burke Neurological Institute and the NIH S10 Shared Instrumentation Grant OD028547-01 for supporting this work. We would like to thank the Center for Neuroanatomy with Neurotropic Viruses, CNNV (P40 OD010996) for sharing the rabies viruses. This work was supported by the National Institute of Neurological Disorders and Stroke (NS100772, NS093002, NS115963 and NS093002).

---

**Figure S1.**
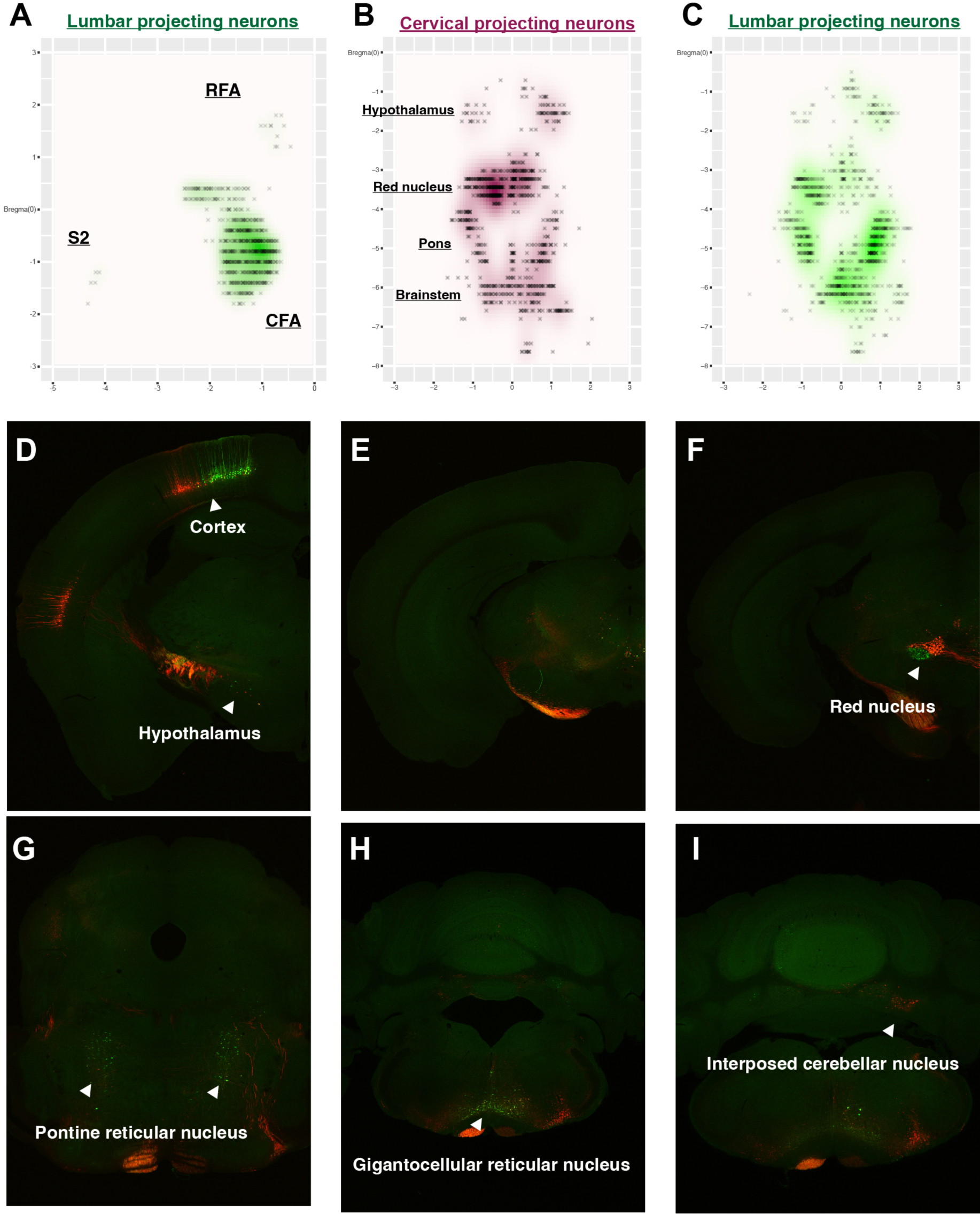

